# Coenzyme Q_4_ is a functional substitute for coenzyme Q_10_ and can be targeted to the mitochondria

**DOI:** 10.1101/2023.07.20.549963

**Authors:** Laura H. Steenberge, Andrew Y. Sung, Jing Fan, David J. Pagliarini

## Abstract

Coenzyme Q_10_ (CoQ_10_) is an important cofactor and antioxidant for numerous cellular processes, and its deficiency has been linked to human disorders including mitochondrial disease, heart failure, Parkinson’s disease, and hypertension. Unfortunately, treatment with exogenous oral CoQ_10_ is often ineffective, likely due to the extreme hydrophobicity and high molecular weight of CoQ_10_. Here, we show that less hydrophobic CoQ species with shorter isoprenoid tails can serve as viable substitutes for CoQ_10_ in human cells. We demonstrate that CoQ_4_ can perform multiple functions of CoQ_10_ in CoQ-deficient cells at markedly lower treatment concentrations, motivating further investigation of CoQ_4_ as a supplement for CoQ_10_ deficiencies. In addition, we describe the synthesis and evaluation of an initial set of compounds designed to target CoQ_4_ selectively to mitochondria using triphenylphosphonium (TPP). Our results indicate that select versions of these compounds can successfully be delivered to mitochondria in a cell model and be cleaved to produce CoQ_4_, laying the groundwork for further development.

## INTRODUCTION

Coenzyme Q (CoQ) is a ubiquitous redox-active lipid with critical functions throughout the cell. It is composed of a polyisoprenoid tail that anchors it in lipid bilayers and a benzoquinone head group that can accept and donate electrons. CoQ is required for shuttling electrons from complexes I and II to complex III in the electron transport chain (ETC) and participates in myriad mitochondrial and extra-mitochondrial pathways including pyrimidine biosynthesis, ferroptosis suppression, fatty acid oxidation, sulfide detoxification, and membrane antioxidation (1). These diverse and important functions are reflected in the wide spectrum of clinical diseases that arise from deficiency of CoQ_10_ (the predominant CoQ species found in humans). Genetic defects in CoQ_10_ biosynthesis cause phenotypes varying from nephropathy and myopathy to fatal multiorgan disease (2). In addition, secondary deficits in CoQ_10_ levels have been observed in aging (3) and in numerous common conditions such as neurodegenerative diseases (4), cardiomyopathy (5), and primary mitochondrial diseases (6).

CoQ_10_ has emerged as an attractive therapeutic candidate to augment CoQ_10_ levels in these disorders. CoQ_10_ treatment has been the subject of numerous clinical trials and is among the most common supplements taken in the Western world (5). Despite widespread interest and use, CoQ_10_ treatment is frequently ineffective (7–9). Poor bioavailability and cell delivery due to the large size and extreme hydrophobicity of CoQ_10_ are often cited as reasons for treatment failure (1, 8, 10). Transportation of the lipophilic CoQ_10_ across the gut epithelium and through the aqueous environment of the body is a major barrier to effective supplementation, with only 2% of orally administered CoQ reaching the bloodstream (11) and an even smaller fraction reaching mitochondria and other target membranes in the cell (12, 13). Thus, a less hydrophobic CoQ analog or an analog better directed to mitochondria could more efficiently reach target membranes and represent attractive alternatives for CoQ_10_ supplementation.

The hydrophobicity of CoQ is primarily determined by its long polyisoprenoid tail (Table 1), which plays a critical role in anchoring CoQ within the core of the lipid bilayer to facilitate its interactions with membrane-embedded proteins. The specific number of isoprene units in the tail (denoted by a subscript) varies between organisms. For example, humans, *Saccharomyces cerevisiae*, and *Escherichia coli* have ten, six, and eight isoprene units in the tail of their dominant CoQ form, respectively. Although the reason for this variation remains poorly understood, it suggests that reducing the hydrophobicity of the CoQ tail may not compromise its function in humans.

**Table 1.**
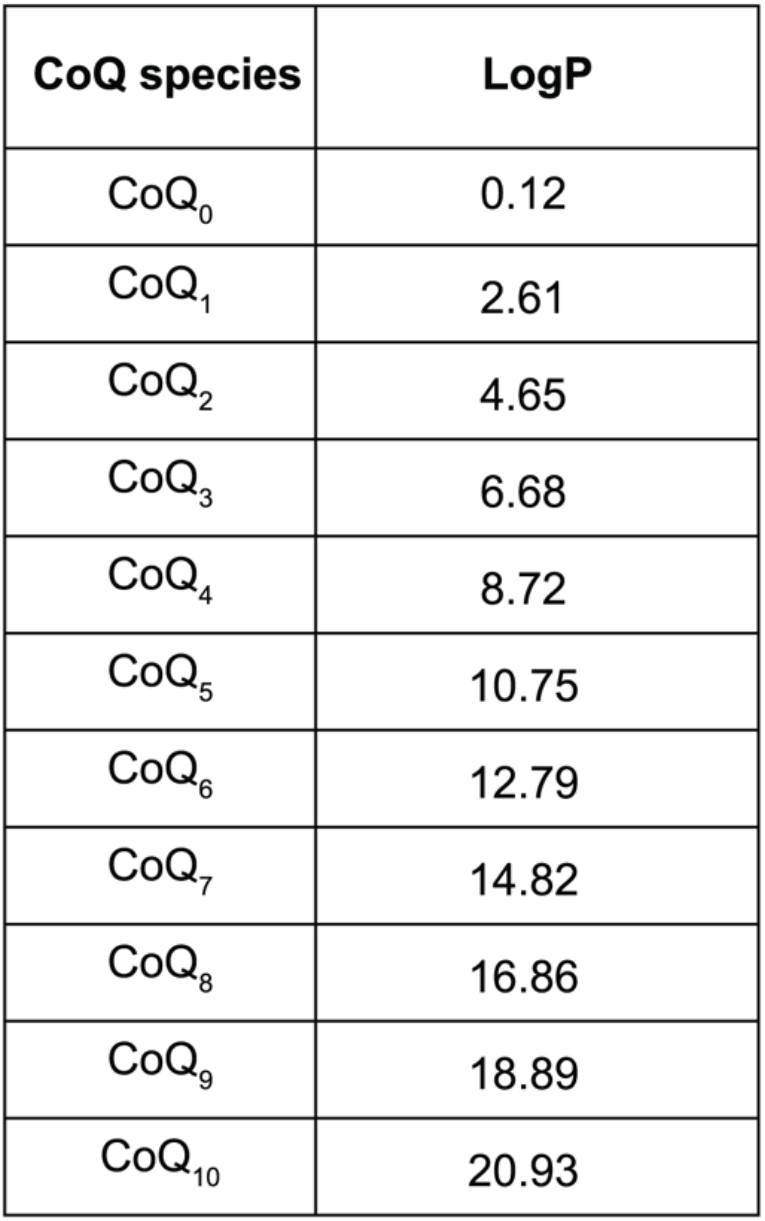
Octanol-water partition coefficients of CoQ species with increasing isoprenoid tail lengths as calculated using Advanced Chemistry Development (ACD) Software.

| CoQ species | LogP |
| --- | --- |
| CoQ<sub>0</sub> | 0.12 |
| CoQ<sub>1</sub> | 2.61 |
| CoQ<sub>2</sub> | 4.65 |
| CoQ<sub>3</sub> | 6.68 |
| CoQ<sub>4</sub> | 8.72 |
| CoQ<sub>5</sub> | 10.75 |
| CoQ<sub>6</sub> | 12.79 |
| CoQ<sub>7</sub> | 14.82 |
| CoQ<sub>8</sub> | 16.86 |
| CoQ<sub>9</sub> | 18.89 |
| CoQ<sub>10</sub> | 20.93 |

CoQ analogs with tail modifications have been investigated as CoQ_10_ substitutes. For instance, idebenone, a CoQ mimetic approved for treatment of Leber hereditary optic neuropathy (14), consists of the benzoquinone head group of CoQ with a fully saturated ten-carbon acyl chain capped by a hydroxyl group in place of the isoprenoid tail. The dramatic decrease in hydrophobicity and structural differences likely prevent its direct substitution for CoQ_10_ in the ETC; rather, its cellular effects occur through entirely separate mechanisms (15, 16). Moreover, studies involving CoQ species with shortened isoprenoid tail lengths in human systems have been met with inconsistent outcomes. In HL-60 cells, CoQ_6_ was unable to increase CoQ-dependent oxidative phosphorylation (OxPhos) activities (17); conversely, a separate study reported that CoQ_4_ restored ATP levels and CoQ-dependent OxPhos in CoQ-deficient patient fibroblasts (18). Additionally, shorter chain quinones have been associated with adverse effects in cells, such as the induction of reactive oxygen species and apoptosis (17, 19–21).

Mitochondria-targeting CoQ species have also been developed as potential CoQ_10_ substitutes. Most notably, the molecule MitoQ is composed of the CoQ head group linked to triphenylphosphonium (TPP), a mitochondriotropic moiety. While MitoQ rapidly accumulates in mitochondria and exhibits potent antioxidant behavior, the attached TPP prevents the head group from properly interacting with ETC complexes (22), and thus it is unable to substitute for CoQ_10_ in supporting OxPhos. Here, toward establishing a more effective CoQ therapeutic, we examined a series of CoQ tail lengths across multiple assays to clarify the minimum hydrophobicity required for CoQ to function in human cells and developed a series of CoQ analogs with reversible TPP linkages to explore their ability to enhance the functional delivery of CoQ.

## RESULTS

### Short-chain CoQ analogs support OxPhos

Although decreasing the tail length of CoQ may improve its bioavailability, its tail must still be sufficiently lipophilic to partition CoQ into the inner leaflet of the mitochondrial inner membrane to interact with ETC complexes. To find this balance between aqueous solubility and functionality, we sought to identify the minimum isoprenoid tail length required for CoQ to support OxPhos. Compromised OxPhos function causes cell death in media containing galactose as the sole carbon source (23). Accordingly, HepG2 *COQ2*^*-/-*^ cells, which lack a key CoQ_10_ biosynthetic enzyme and are CoQ_10_-deficient, had increased rates of cell death in galactose media, which were reduced with 20 µM CoQ_10_ supplementation (Figure 1a). Surprisingly, CoQ_4_ supplementation at this concentration was equally effective at preventing cell death. CoQ_2_ supplementation initially maintained viability in galactose (Supporting Information Figure S1a), but extended treatment caused cell death, likely due to inherent toxicity of the compound (20, 21) (Supporting Information Figure S1b). CoQ_4_ showed no evidence of cellular toxicity or increased ROS production at concentrations up to 100 µM (Supporting Information Figure S2).

**Figure 1.**
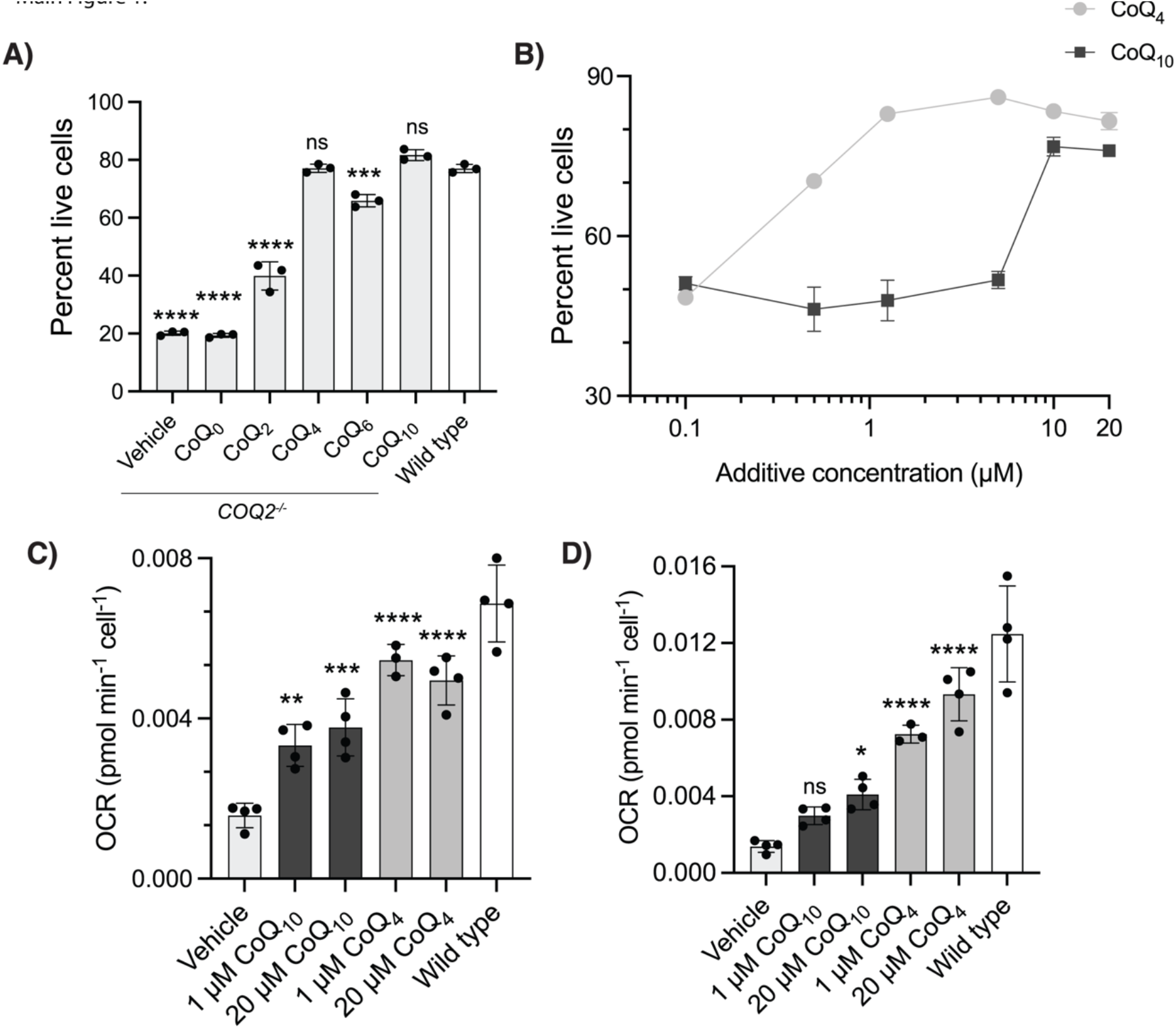
Short-chain CoQ analogs support OxPhos. **a** Percent live wild-type or *COQ2*^*-/-*^ HepG2 cells after 72 hours of incubation in galactose media with 20 µM indicated CoQ additive. Cells were pretreated with CoQ additives for 48 hours to ensure sufficient uptake. Live cells defined as not staining for AAD-7 or annexin-V. *n* = three independent technical replicates, error bars indicate standard deviation. **b** Survival of *COQ2*^*-/-*^ HepG2 cells in galactose as in **a** over a titration of CoQ_4_ and CoQ_10_ concentrations. *n* = two independent technical replicates, error bars indicate standard deviation. **c-d** Oxygen consumption rates (OCR) indicating **c** basal respiration and **d** maximal respiration in *COQ2*^*-/-*^ and wild-type HepG2 cells after 24 hour incubation with CoQ additives. *n* = four independent technical replicates, error bars indicate standard deviation. *ns: not significant, * p < 0*.*05, ** p < 0*.*01, *** p < 0*.*001, *** p < 0*.*0001 For exact p values see Supporting Information Table S1*.

We repeated this analysis across a range of concentrations using CoQ_4_ and CoQ_10_ and found that, comparatively, ten-fold lower concentrations of the former were needed to rescue cell viability (Figure 1b). Importantly, we confirmed that CoQ_4_ supplementation also increased basal and maximal respiration in CoQ_10_-deficient cells to a greater extent than CoQ_10_ supplementation (Figure 1c, 1d). Overall, these results indicate that a tail length of four isoprene units is sufficient to support proper OxPhos function in human cells. Interestingly, while 1 µM of CoQ_10_ was not sufficient to rescue viability in galactose, it was able to increase basal OCR to a similar level as 20 µM CoQ_10_, suggesting that other functions of CoQ might contribute to viability in galactose media.

### CoQ_4_ can support ferroptosis defense and pyrimidine biosynthesis

Next, we examined whether CoQ_4_ could participate in CoQ_10_ roles beyond the ETC, such as ferroptosis defense and *de novo* pyrimidine biosynthesis. Reduced CoQ_10_ (CoQ_10_H_2_) at the plasma membrane helps prevent ferroptosis through radical trapping and suppression of lipid peroxidation and is regenerated by the oxidoreductase FSP1 for continued ferroptosis suppression (24, 25). This pathway works in parallel with another major ferroptosis defense mechanism mediated by GPX4, a hydroperoxidase that neutralizes lipid peroxides using glutathione. Thus, the HepG2 *COQ2*^*-/-*^ cells, which lack CoQ_10_ at the plasma membrane, were more sensitive to the GPX4 inhibitor RSL3 (Figure 2a). This RSL3 sensitivity was decreased by CoQ_10_ supplementation and completely ablated by CoQ_4_ (Figure 2b). Inhibition of FSP1 by its inhibitor iFSP1 blocked the ability of both CoQ_10_ and CoQ_4_ (at low concentrations) to prevent ferroptosis (Figure 2c). Higher CoQ_4_ supplementation concentrations caused resistance to FSP1 inhibition, likely because there were sufficient amounts of reduced CoQ_4_ at the plasma membrane to preclude the need for FSP1-mediated regeneration.

**Figure 2.**
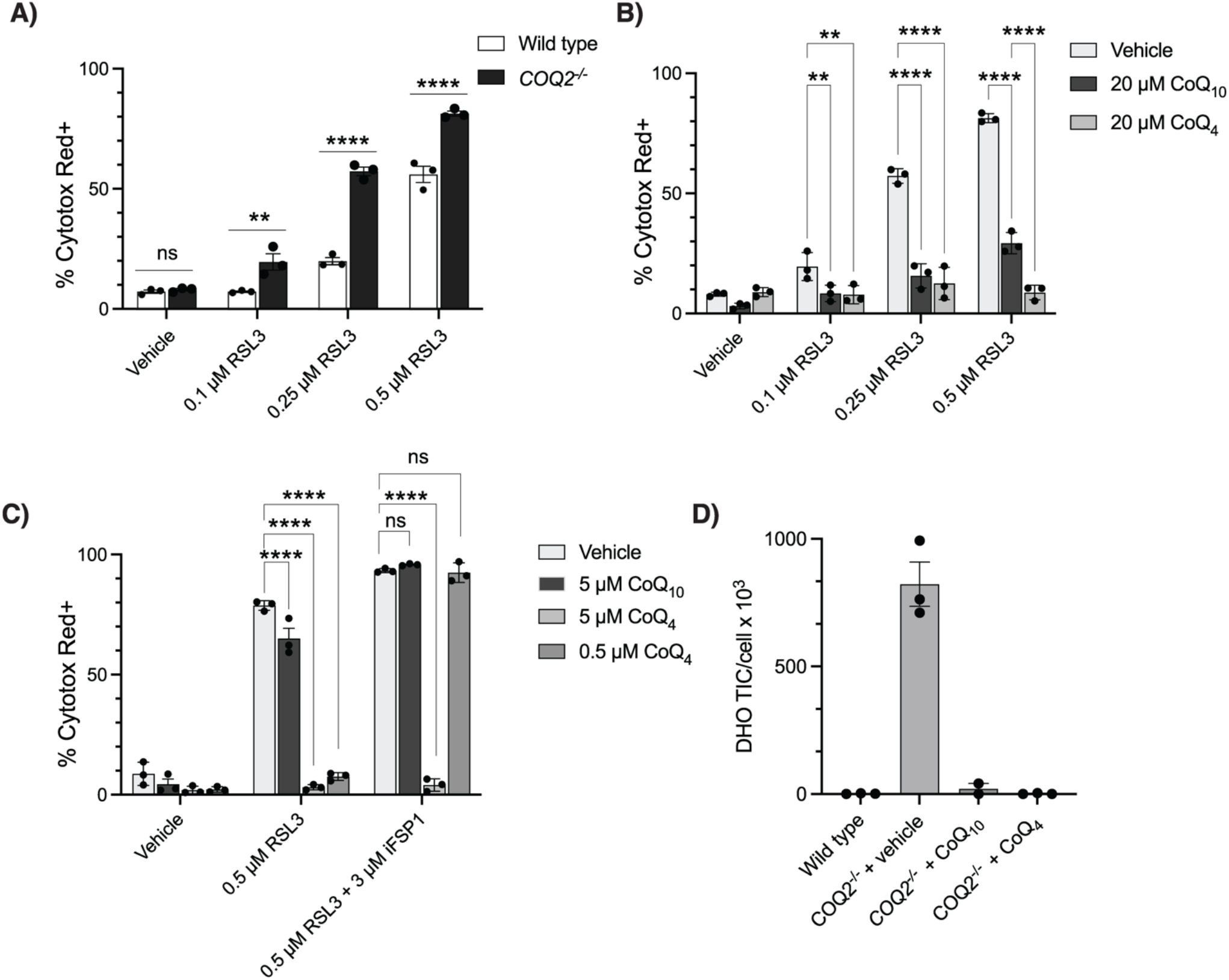
CoQ_4_ can participate in ferroptosis suppression and pyrimidine biosynthesis. **a** Cell death of wild-type and *COQ2*^*-/-*^ HepG2 cells after 24 hours of treatment with GPX4-inhibitor RSL3. Cell death determined by high levels of Cytotox Red labeling. *n* = three independent technical replicates, error bars indicate standard error. **b** Cell death of *COQ2*^*-/-*^ HepG2 cells after 24 hours of treatment with RSL3 following 24 hours of pre-incubation with 20 µM CoQ_4_ or CoQ_10_. *n* = three independent technical replicates, error bars indicate standard error. **c**. Cell death of *COQ2*^*-/-*^ HepG2 cells after 24 hours of treatment with RSL3 or iFSP1 following 24 hours of pre-incubation with indicated concentrations of CoQ_4_ or CoQ_10_. *n* = three independent technical replicates, error bars indicate standard error. **d** Total ion counts of dihydroorotate normalized to cell count in HAP1 wild-type and *COQ2*^*-/-*^ cells after 24 hours of treatment with 20 µM CoQ_4_ or CoQ_10_. *n* = three independent technical replicates, error bars indicate standard error. *ns: not significant, * p < 0*.*05, ** p < 0*.*01, *** p < 0*.*001, *** p < 0*.*0001 For exact p values see Supporting Information Table S1*.

CoQ also plays a key role in the pyrimidine *de novo* biosynthetic pathway as a cofactor for dihydroorotate dehydrogenase (DHODH); thus, HAP1 *COQ2*^*-/-*^ cells accumulated dihydroorotate (DHO), the substrate of DHODH (Figure 2d). Supplementation of both CoQ_4_ and CoQ_10_ restored DHO to wild-type levels (Figure 2d), suggesting that DHODH can use CoQ_4_ to synthesize orotate. Altogether, these results indicate that CoQ_4_ can fulfill roles of CoQ_10_ outside of the ETC.

### CoQ_4_ can be preferentially enriched in mitochondria

While CoQ_4_ is much less hydrophobic than CoQ_10_, it still contains 20 aliphatic carbons in its tail and retains considerable lipophilicity (Table 1). Therefore, there will likely still be significant barriers to its bioavailability and cellular delivery. We hypothesized that a reversible linkage to triphenylphosphonium (TPP), a mitochondria-targeting group, could increase the delivery of functional CoQ_4_. TPP is a well-studied lipophilic cation known to selectively transport cargo into mitochondria, driving concentrations of cargo in the mitochondrial matrix 100- to 1000-fold higher than extracellular concentrations. Moreover, TPP-conjugated compounds often bypass traditional transport mechanisms, instead directly passing through membranes to reach the mitochondrial matrix (26). Exogenous CoQ taken up by the cell primarily becomes trapped in lysosomes during transport (12, 13); thus, bypassing the cellular transport mechanisms with TPP could increase the amount of CoQ reaching target membranes.

Studies with MitoQ have demonstrated that the irreversible attachment of TPP to CoQ prevents proper interactions with complex I and complex III, likely due to steric hindrance and improper membrane portioning. Therefore, we attached the TPP moiety to the head group of CoQ through an ester linkage (Figure 3a). Esters are often labile in cells and can be hydrolyzed by resident nonspecific esterases (27, 28). We synthesized three CoQ_4_-TPP compounds with varying acyl linker chain lengths (Figure 3b). Porcine liver esterase (PLE), a representative esterase, hydrolyzed all three compounds to a large extent *in vitro* (Figure 4a) while a similar CoQ_10_-TPP compound was unable to be hydrolyzed (Figure 4b). Incubating cells with CoQ_4_-TPP resulted in the dose-dependent production of free CoQ_4_ (Figure 4c), although the resulting levels were ten-fold lower than levels from treatment with exogenous unmodified CoQ_4_ (Figure 4d). Importantly, while unmodified CoQ_4_ treatment resulted in comparable amounts of CoQ_4_ in the cytoplasmic and mitochondrial fractions of the cell, CoQ_4_-TPP increased relative CoQ_4_ levels in the mitochondrial fraction compared to the cytoplasmic fraction (Figure 4e-g, Supporting Information Figure S3). Unfortunately, further investigations of CoQ_4_-TPP were limited by the toxicity of the compound (Supporting Information Figure S4). While optimization of the delivery system to reduce toxicity and improve hydrolysis is required, these data provide proof-of-concept that CoQ_4_ can be reversibly linked to a mitochondria-targeting group and preferentially delivered to mitochondria, providing a platform for further study into CoQ targeting strategies.

**Figure 3.**
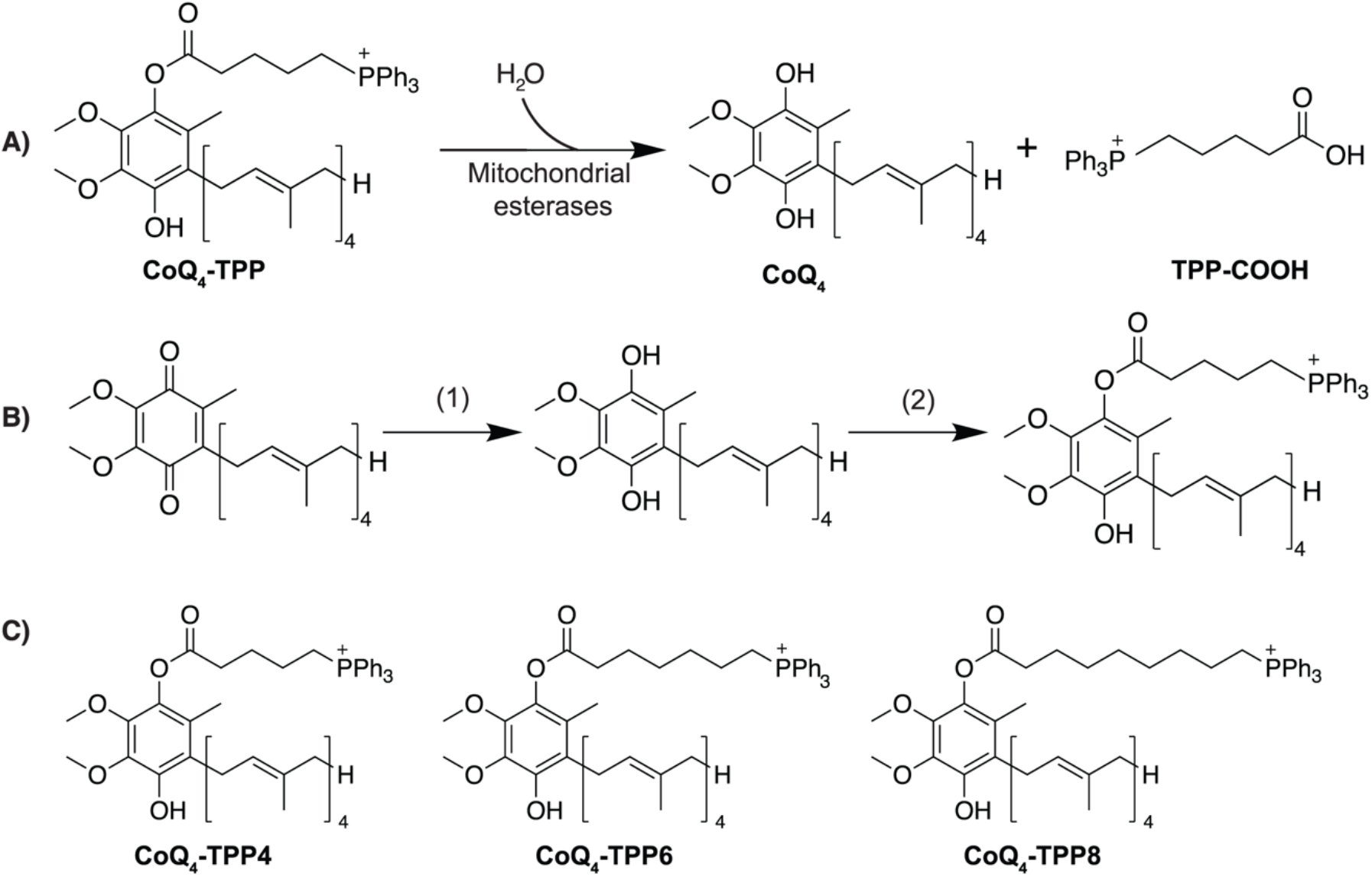
Schematic of CoQ_4_-TPP hydrolysis and synthetic scheme. **a** Schematic of CoQ_4_-TPP hydrolysis by mitochondrial esterases to produce CoQ_4_ and a carboxy-TPP side product (TPP-COOH). **b**. Chemical synthesis of CoQ_4_-TPP. Reaction conditions: (1) NaBH_4_ in methanol/isopropanol. (2) 4-carboxybutyl triphenylphosphonium bromide, EDC, DMAP in dichloromethane. **c**. Structures of CoQ_4_-TPP compounds synthesized for this study.

**Figure 4.**
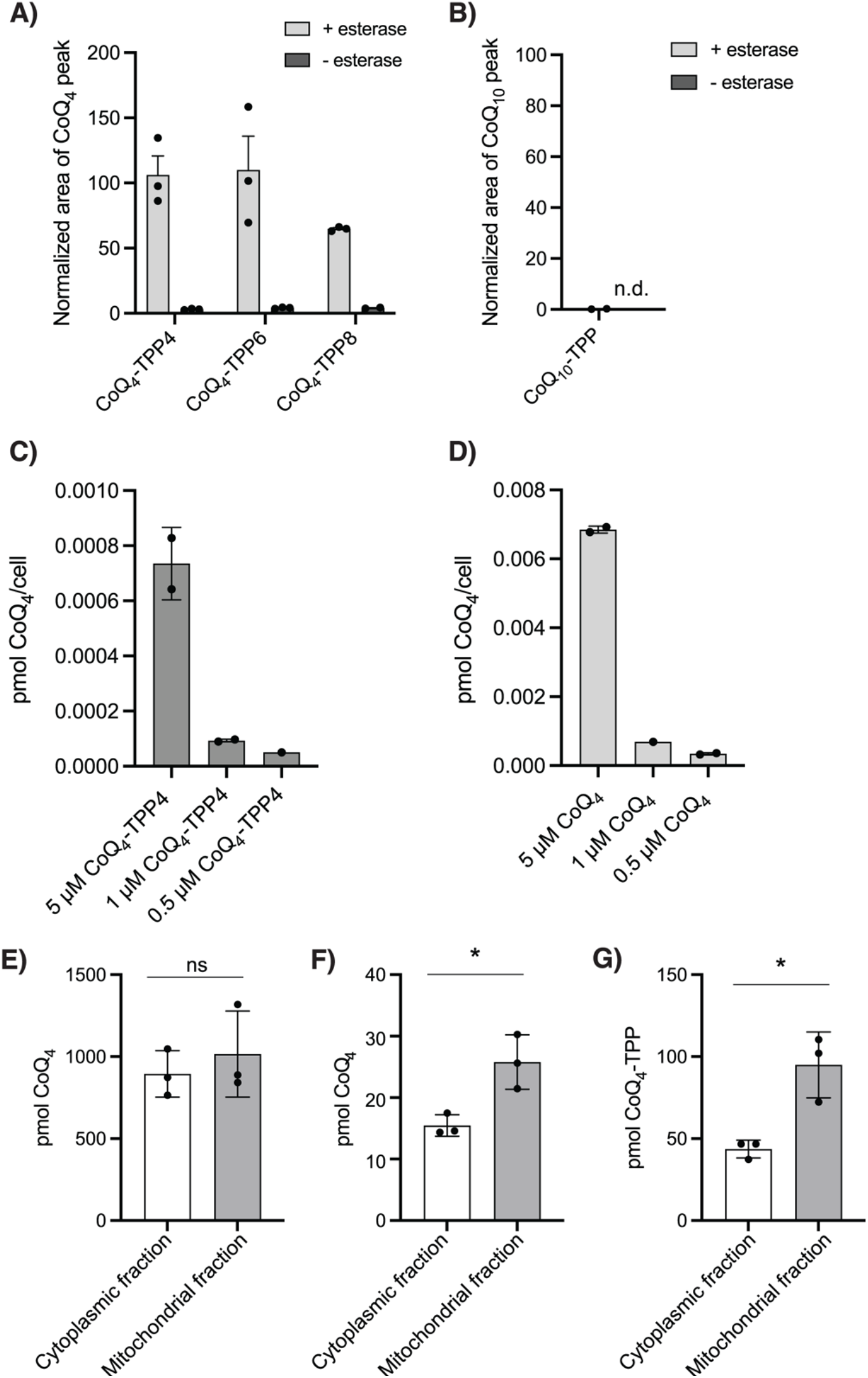
CoQ_4_-TPP can be hydrolyzed to produce CoQ_4_. **a-b** CoQ production from CoQ-TPP species after 24 hours of incubation with porcine liver esterase, normalized to equivalent CoQ_4_ (**a**) or CoQ_10_ (**b**) area indicating 100% turnover. CoQ levels measured by HPLC-ECD. *n* = three independent technical replicates, error bars indicate standard error. **c** Levels of CoQ_4_ after 24 hours of incubation of wild-type HepG2 cells with CoQ_4_-TPP (**c**) or unmodified CoQ_4_ (**d**) measured by HPLC-ECD after cellular lipid extraction. *n* = two independent technical replicates, error bars indicate standard deviation. **e – f**. CoQ_4_ levels in different subcellular compartments after 3 hours of incubation of HepG2 cells with 10 µM CoQ_4_ (**e**) or CoQ_4_-TPP4 (**f**). **g**. CoQ_4_-TPP levels in different subcellular compartments after 3 hours of incubation of HepG2 cells with 10 µM CoQ_4_-TPP4. *n* = three independent technical replicates, error bars indicate standard error. *ns: not significant, * p < 0*.*05, ** p < 0*.*01, *** p < 0*.*001, *** p < 0*.*0001 For exact p values see Supporting Information Table S1*.

## DISCUSSION

Stemming from its myriad critical cellular roles, CoQ_10_ deficiency leads to a variety of human diseases. CoQ_10_ supplementation is often recommended for numerous conditions including primary mitochondrial diseases, heart failure, Parkinson’s disease, and hypertension (5, 29). CoQ_10_ is widely used as a nutritional supplement, with a global market size estimated to be nearly 600 million USD per year (8). Unfortunately, the evidence supporting its clinical efficacy is often weak and treatment failure is common, likely because the extreme hydrophobicity and large size of CoQ_10_ limits its uptake and distribution to target membranes. Here, we demonstrate that the less hydrophobic CoQ_4_ can fulfill multiple functions of CoQ_10_ in a cellular model, positing that it could be a viable therapeutic alternative for CoQ supplementation, and show that CoQ_4_ can be preferentially targeted to mitochondria through linkage to TPP.

Surprisingly, we found that CoQ_4_ could rescue CoQ-deficiency phenotypes in cells at much lower treatment concentrations than CoQ_10_. This likely reflects increased CoQ_4_ delivery to target membranes rather than enhanced CoQ_4_ functionality. Indeed, in a reconstituted system, complex I displayed decreased catalytic efficacy with shorter CoQ species including CoQ_4_ (30). CoQ_4_ must reach multiple cellular membranes to fulfill CoQ’s various cellular roles. Its movement throughout the cell could be mediated by the endomembrane system, as has been shown for CoQ_10_ (31), or could occur through a separate mechanism. Given that CoQ_4_ has been shown to exchange between phospholipid bilayers of two vesicle populations while CoQ_10_ cannot (32), it is possible that CoQ_4_ does not require the same level of active transport between membranes as CoQ_10_ but instead can move through the cell without relying on dedicated protein machinery.

Assessing the bioavailability and cellular delivery of CoQ_4_ in an animal model will provide further insight into its potential as a CoQ_10_ substitute. While CoQ_4_ is much less hydrophobic than CoQ_10_, it still possesses a highly hydrophobic 20-carbon tail and could experience similar bioavailability barriers as CoQ_10_. Idebenone, a CoQ analog with a fully saturated 10-carbon tail, is much less lipophilic than CoQ_10_ yet is still poorly water soluble, which presents many challenges to its use (33–35). Thus, additional strategies to boost CoQ_4_ delivery will likely be required to develop an effective therapeutic.

Toward further improving CoQ delivery, we trialed reversibly attaching TPP to the CoQ head group, as TPP can drive cargo molecules to high concentrations intracellularly and specifically in mitochondria. Encouragingly, we observed release of free CoQ_4_ that was enriched in mitochondria. Unexpectedly, there was a substantial amount of CoQ_4_ and CoQ_4_-TPP in the cytoplasmic fraction as well. We suspect that this discrepancy from the usual profound enrichment of TPP compounds in mitochondria is due to the known disruption of the inner mitochondrial membrane potential caused by hydrophobic TPP molecules (36–38), which would decrease the electrochemical gradient driving CoQ_4_-TPP into mitochondria and likely also contribute to the observed toxicity. A less disruptive mitochondria targeting group, such as the novel TPP-CF_3_ derivatives (37), could potentially target CoQ_4_ more effectively. Additionally, we surmise that the toxicity of our compound arises from the unhydrolyzed CoQ_4_-TPP, as joint treatment with unmodified CoQ_4_ and TPP-COOH resulted in high intracellular levels without evident toxicity. Therefore, more labile linker constructs would likely decrease the toxicity of the compound while also leading to higher levels of CoQ_4_ enrichment. Overall, our work encourages further exploration of less hydrophobic CoQ analogs in therapeutic regimens and provides proof-of-concept that CoQ species can be reversibly linked to a targeting moiety to enhance delivery to mitochondria.

### EXPERIMENTAL PROCEDURES

#### General cell culture

HepG2 lines (ATCC) cultured in DMEM (Thermo) supplemented with 10% heat-inactivated FBS (R&D Systems, S11550) and 1x penicillin-streptomycin (Pen/Strep; Thermo, 15140122) at 37°C and 5% CO_2_. The Genome Engineering and iPSC Center (GEiC) at Washington University in St. Louis generated the HepG2 *COQ2*^*-/-*^ cell lines. HAP1 lines (Horizon Discovery) cultured in IMDM supplemented with 10% heat-inactivated FBS and 1x penicillin-streptomycin at 37°C and 5% CO_2_. Lines regularly tested for mycoplasma contamination. Cell counts obtained using a Muse Cell Analyzer (Luminex) and the Muse Count and Viability Assay (Luminex).

#### Galactose Rescue

Wild-type and *COQ2*^-/-^ HepG2 cells were plated at 75,000 cells/well in 6-well plates. After allowing cells to adhere overnight, medium was exchanged to DMEM containing either vehicle (isopropanol) or CoQ additives in isopropanol. After 48 hours of incubation, cells were washed with dPBS and medium was replaced with glucose-free DMEM containing 10 mM of galactose, 1 mM pyruvate, and 100 µg/mL uridine along with CoQ additives or vehicle. After 72 hours, cell apoptosis was assessed by the Muse Annexin-V and Dead Cell Kit. For shorter CoQ_2_ supplementation, cells were incubated with 20µM CoQ_4_ for 24 hours prior to galactose switch, and apoptosis was assessed after 48 hours in galactose.

#### Seahorse Measurements

An Agilent Seahorse XFe96 Analyzer was used to determine oxygen consumption rates (OCR). Cells were plated at 30K cells per well in XF96 microplates (Agilent, 102416-100) with media containing CoQ species. The next day, cell media was exchanged for XF assay medium supplemented with 25 mM glucose, 1 mM pyruvate, 4 mM glutamine (Agilent, 03680-100), and CoQ species (Sigma). Oxygen consumption rate was measured at baseline and after injections of 2 µM oligomycin, 3 µM FCCP, and 0.5 µM rotenone/antimycin A (Agilent, 103015-100). Rates were normalized to cell count per well obtained immediately prior to XF assay medium exchange, obtained by Sartorius IncuCyte S3 live-cells analysis system.

#### Ferroptosis Sensitivity

Wild-type and *COQ2*^-/-^ HepG2 cells were plated at 25,000 cells/well in a 24 well plate and allowed to adhere overnight. Medium was exchanged to DMEM with 1 mM pyruvate and 100 µg/mL uridine along with either vehicle (isopropanol) or CoQ additives (Sigma) in isopropanol. After 48 hours of incubation, cells were washed with dPBS. Media was replaced with DMEM with 1 mM pyruvate, 100 µg/mL uridine, and Incucyte Cytotox Red Reagent (Fisher, NC1015259); along with RSL3 (Sigma, SML2234), iFSP1 (Fisher, NC1755669) or vehicle (DMSO). Cell death was quantified after 24 hours using a Sartorius IncuCyte S3 live-cells analysis system.

#### *DCFDA and Cell* Growth *Assay*

Cells were plated at 40,000 cells/well in a 24 well plate and allowed to adhere overnight. Cell media was exchanged to DMEM containing 5 µM DCFDA (Abcam, ab113851) and incubated for 30 minutes at 37°C. Cells were washed twice with dPBS and medium was replaced with DMEM containing the CoQ additives (Sigma), menadione (Sigma, M5625), or vehicle (isopropanol). Using a Sartorius IncuCyte S3 live-cells analysis system, green fluorescence was determined to obtained ROS measurements and cell number was obtained to calculate fold change.

#### Dihydroorotate measurements

HAP1 wild-type and *COQ*2^*-/-*^ cells were plated at 500,000 cells/well in a 6 well plate and allowed to adhere overnight. Media was exchanged to DMEM containing either vehicle, 20 µM CoQ_4_ (Sigma, C2470) or 20 µM CoQ_10_ (Sigma, C9538) in isopropanol. After 24 hours of incubation, cells were washed 3 times with cold dPBS, then incubated at -80°C with cold LC-MS grade 80:20 methanol/water (v/v) for 15 minutes. Cells collected and centrifuged at 16,000 xg for 5 minutes. Supernatant collected and dried under nitrogen flow. To measure dihydroorotate, samples resuspended in LC-MS grade water. Samples were analyzed using a Thermo Q-Exactive mass spectrometer coupled to a Vanquish Horizon UHPLC. Analytes were separated on a 100 × 2.1 mm, 1.7 µM Acquity UPLC BEH C18 Column (Waters), with a 0.2 ml min^−1^ flow rate and with a gradient of solvent A (97:3 H_2_O/methanol, 10 mM TBA, 9 mM acetate, pH 8.2) and solvent B (100% methanol). The gradient is: 0 min, 5% B; 2.5 min, 5% B; 17 min, 95% B; 21 min, 95% B; 21.5 min, 5% B. Data were collected in full-scan negative mode. Setting for the ion source were: 10 aux gas flow rate, 35 sheath gas flow rate, 2 sweep gas flow rate, 3.2 kV spray voltage, 320°C capillary temperature and 300°C heater temperature. The metabolites reported here were identified based on exact *m*/*z* and retention times determined with chemical standards.

#### Synthesis of CoQ-TPP Compounds

To synthesize CoQ_4_-TPP compounds, CoQ_4_ (100mg, 2.26 µmol) was added to isopropanol/methanol (10 mL). CoQ_4_ for synthesis obtained from WuXi Chemicals. NaBH_4_ (Sigma, 480886) added to CoQ until color changed from orange to colorless was observed, indicating reduction. Reaction quenched with 1M hydrochloric acid (20 mL), diluted with dichloromethane (15 mL), and washed with 1M hydrochloric acid (50 mL) then brine (50 mL). Organic layer sealed under argon. 4-dimethylaminopyridine (25 mg, 2.0 µmol; Sigma, 107700), N-(3-dimethylaminopropyl)-N’ethylcarbodiimide hydrochloride (45 mg, 2.0 µmol; Sigmal, 03450), and 4-carboxybutyl- (Sigma, 157945); 6-carboxyhexyl (WuXi Chemicals) or 8-carboxyoctyl (Toronto Research Chemicals, C178905) triphenylphosphonium bromide (100 mg, 2.0 µmol) dissolved in dichloromethane and added dropwise to reduced CoQ. Reaction stirred at room temperature for 72 hours, and product formation confirmed by TLC. Solvent removed under reduced pressure, and column chromatography on silica gel with 2:1 ethyl acetate:methanol gave products as orange gel. Remaining silica gel filtered with acetonitrile elution. Yield calculated at 5-10% for monoacylated products.

CoQ_4_-TPP_4_: ^1^H NMR 7.87 (t, 6H), 7.77 (t, 3H), 7.68 (t, 6H), 5.09 (m, 4H), 4.00 (m, 2H), 3.87 (m, 2H), 3.67 (s, 2H), 3.65 (s, 2H), 3.32 (d, 1H), 3.09 (d, 1H), 2.11 (d, 3H), 2.05 (d, 5H, 1.97 (d, 5H), 1.68 (s, 9H), 1.60 (m, 12H), 1.23 (s, 6H), 0.89 (m, 3H). ^13^C NMR 134.89, 133.85, 130.49, 124.42, 118.92, 118.24, 96.15, 60.83, 39.75, 32.94, 29.72, 26.79, 25.72, 25.42, 21.88, 20.39, 17.72, 16.03, 12.12. HRMS (ESI-MS, m/z) calculated 801.46, found 801.465.

CoQ_4_-TPP_6_: ^1^H NMR 7.89 (t, 6H), 7.78 (t, 3H), 7.70 (t, 6H), 5.09 (m, 4H), 3.97 (m, 2H), 3.88 (m, 2H), 3.76 (s, 2H), 3.33 (d, 1H), 3.14 (d, 1H), 2.56 (M, 2H), 2.12 (s, 2H), 2.05 (d, 5H, 1.97 (d, 6H), 1.76 (s, 15H), 1.68 (m, 9H), 1.60 (s, 9H), 1.46 (m, 3H), 0.93 (m, 2H). ^13^C NMR 172.11, 144.87, 137.64, 134.89, 133.75, 130.4, 124.43, 118.99, 118.3, 96.15, 60.98, 39.75, 33.72, 29.81, 28.53, 26.80, 25.73, 24.5, 22.45, 17.72, 16.02, 12.12. HRMS (ESI-MS, m/z) calculated 829.50, found 829.495.

CoQ_4_-TPP_8_: ^1^H NMR 7.86 (t, 6H), 7.79 (t, 3H), 7.71 (t, 6H), 5.10 (m, 4H), 4.0 (m, 2H), 3.9 (m, 3H), 3.79 (m, 2H), 3.7 (m, 1H), 3.34 (d, 1H), 3.2 (d, 1H), 2.55 (m, 1H), 2.48 (m, 1H), 2.05 (m, 6H), 2.0 (m, 6H), 1.81 (m, 18H), 1.68 (s, 6H), 1.60 (m, 12H), 1.26 (m, 9H), 0.89 (m, 2H). ^13^C NMR 135.99, 134.87, 133.74, 130.50, 130.40, 124.42, 119.02, 118.34, 96.14, 39.75, 29.73, 28.83, 27.36, 26.79, 25.73, 24.99, 24.15, 22.72, 17.72, 16.05. HRMS (ESI-MS, m/z) calculated 857.53, found 857.526.

#### HPLC-ECD Measurement of CoQ levels

Samples were injected into HPLC (Ultimate 3000, Thermo Scientific) with an electrochemical detector (ECD-3000RS). The first electrode (6020RS) was set to +600mV and placed before the column (Thermo Scientific, Betasil C18, 100 × 2.1 mm, 3 µM particle) to oxidize all quinones. The second electrode (6011RS) was set to - 600mV to reduce all quinones exiting the column, and the third electrode was set at +600mV to measure redox active species. Peaks were quantified with Chromeleon 7.2.10 software. To measure CoQ_4_ levels, extracted lipids were resuspended in methanol, and the mobile phase was 95% methanol 5% 1M ammonium acetate pH 4.4 in water. To measure CoQ_10_ levels, extracted lipids were resuspended in isopropanol, and the mobile phase was 78% methanol, 20% isopropanol, and 2% 1M ammonium acetate pH 4.4 in water.

#### Representative Esterase Hydrolysis

CoQ-TPP compounds were incubated at 200 µM at 37°C for 24 hours in KCl buffer (120 mM KCl, 10 mM HEPES, 1 mM EGTA, pH = 7.2) containing either 1 mg/mL porcine liver esterase or a buffer control along with CoQ_2_ (Sigma, C8081) as an internal standard. To extract CoQ_4_, cold LC-MS grade methanol was added and sample was incubated at -80°C for 15 minutes. Sample vortexed for 5 minutes at 4°C in a disruptor genie set to max (3000 rpm) and centrifuged at 16,000 xg for 5 minutes. Supernatant collected, dried under nitrogen flow, and CoQ species measured by HPLC-ECD. To extract CoQ_10_, 400µL petroleum ether added to reaction. Samples vortexed for 10 minutes at 4°C in a disruptor genie set to max (3000 rpm) then centrifuged at 1000 xg for 3 minutes. Petroleum ether layer collected and 400 µL fresh petroleum ether added to reaction mixture. Samples vortexed for 3 minutes at 4°C in a disruptor genie set to max (3000 rpm) then centrifuged at 1000 xg for 3 minutes. Petroleum ether layer collected and combined with prior petroleum ether, dried under nitrogen flow, and CoQ species measured by HPLC-ECD. For hydrolysis prior to cell incubation, reactions prepared similarly. After overnight incubation, 100µL sample was added to DMEM for final concentration of 5µM of CoQ-TPP species. CoQ_4_ was extracted as above and measured via HPLC-ECD.

#### Extraction of CoQ species

Wild-type HepG2 cells wells were seeded at 3,000,000 cells/well in a 6 well plate and allowed to adhere overnight. Medium was exchanged to DMEM containing either vehicle (isopropanol), CoQ_4_ (Sigma, C2470), or CoQ_4_-TPP. After 24 hours, cells were trypsinized and pelleted (600 xg, 5 minutes). Pellet was washed with DBPS + 10% isopropanol, then DBPS + 30% isopropanol. For CoQ_4_, pellet was flash frozen in 500 µL cold methanol with 2 µM of CoQ_6_ (Avanti, 900150O) as an internal standard. Cells were lysed by vortexing for 10 minutes at 4°C in a disruptor genie set to max (3000 rpm) speed. Sample was spun 5 minutes 16,000 xg and the supernatant was collected. 500µL of cold methanol was added and sample was vortexed for 3 minutes and then spun as above. Supernatant was again collected and combined with supernatant from previous step and dried under nitrogen gas.

#### Mitochondria Subfractionation and CoQ extraction

HepG2 cells were seeded in 15cm plates and allowed to adhere overnight. Media was exchanged for DMEM with 10µM either CoQ_4_-TPP or CoQ_4_ (Sigma, C2470). After three hours, mitochondrial fractions were isolated from cells through differential centrifugation as described previously (Frezza, 2007). Briefly, cells and media were collected and washed with dPBS then dDPBS + 10% isopropanol. Cells were resuspended in isolation buffer (10 mM Tris-MOPS, pH 7.4, 1 mM EGTA/Tris, 200 mM sucrose) and homogenized with 50 strokes of a glass-Teflon potter. Whole cell and nuclei were pelleted (10 min x 600g, 4°C), supernatant collected and centrifuged to produce mitochondrial fraction (10 min x 7,000g, 4°C). Supernatant of hard spin collected as cytoplasmic fraction. Protein content in fractions was quantified using Pierce BCA Protein Assay Kit. (Thermo, 23225). 50 ug protein from each fraction saved for later analysis by western blot.

To extract CoQ_4_, the mitochondrial fraction was resuspended in an equivalent amount of isolation buffer to cytoplasmic fraction. To extract CoQ_4_, 400µL of chloroform with CoQ_6_ (Avanti, 900150O) as an internal standard was added and the sample vortexed for 10 minutes at 4°C in a disruptor genie set to max (3000 rpm) speed. Chloroform was collected and process repeated. Lipids were dried and measured with HPLC-ECD.

To assess fraction purity, the 50 ug protein collected was methanol precipitated. Sample was solubilized in RIPA buffer and 1mL methanol added. Sample was pelleted (20 min, 16000xg) and supernatant discarded. 1 mL 90:10 methanol:water (v/v) was added, sample was pelleted (20 min, 16000xg) and supernatant discarded. Precipitated protein was resuspended in sample buffer (Invitrogen, NP0007) to a final concentration of 1 mg/mL. 10 μg of protein from each sample was separated on a NuPAGE 10% Bis-Tris gel (Thermo, NP0303BOX) with a protein standard (Licor, 928-60000). Resolved proteins were then transferred to PVDF membrane (Fisher, IPFL00010), probed with primary antibodies (Abcam, 154856; Cell Signaling, 9644S) followed by HRP-conjugated secondary antibody (Cell Signaling Technologies, 7074S, 7076S) and ECL substrate (Thermo, 34579, 34094). Blots were imaged and analyzed using Azure5 Imaging System (version 1.9.0.0406).

#### CoQ-TPP Toxicity Cell Counts

Wild-type HepG2 cells seeded at 150,000 cells/well in 6 well plate and allowed to adhere overnight. Media was exchanged for DMEM containing CoQ-TPP additives and incubated for 24 hours. Cells were washed with dPBS, collected, and counted using a Muse Cell Analyzer and the Muse Count and Viability Assay (Luminex).

## DATA AVAILABILITY

All data contained within the manuscript.

## SUPPORTING INFORMATION

This article contains supporting information.

## ACKNOWLEDGEMENTS

We thank members of the Pagliarini and Fan labs for helpful discussions throughout this project.

## FUNDING AND ADDITIONAL INFORMATION

This work was supported by NIH awards R35GM131795 (D.J.P.), T32GM008505 (L.H.S), T32GM140935 (L.H.S. and A.Y.S.), T32AG000213 (A.Y.S.) and funds from the BJC Investigator Program (D.J.P.). This study made use of the National Magnetic Resonance Facility at Madison, which is supported by NIH grant P41GM136463, P41GM103399 (NIGMS). Equipment was purchased with funds from the University of Wisconsin-Madison, the NIH P41GM136463, P41GM103399, S10RR02781, S10RR08438, S10RR023438, S10RR025062, S10RR029220), the NSF (DMB-8415048, OIA-9977486, BIR-9214394), and the USDA. We thank P. Cobra for assistance with NMR data acquisition. This study made use of the Washington University in St. Louis Genome Engineering and iPSC Center for cell line generation. The content is solely the responsibility of the authors and does not necessarily represent the official views of the National Institutes of Health.

## ABBREVIATIONS AND NOMENCLATURE

The abbreviations used in this study are:

ATP: adenosine triphosphate
BCA: bicinchoninic acid
CoQ: coenzyme Q
DCFDA: dichlorodihydrofluorescein diacetate
DHO: dihydroorotate
DHODH: dihydrooratate dehydrogenase
DMEM: Dulbecco’s Modified Eagle Media
dPBS: Dulbecco’s phosphate buffered saline
ETC: electron transport chain
FBS: fetal bovine serum
FCCP: Carbonyl cyanide p-trifluoro-methoxyphenyl hydrazone
HEPES: 4-(2-hydroxyethyl)-1-piperazineethanesulfonic acid
HPLC-ECD: high performance liquid chromatography – electrochemical detection,
IMDM: Iscove’s Modified Dulbecco’s Medium
KCl: potassium chloride,
OxPhos: oxidative phosphorylation
PLE: porcine liver esterase
RIPA: radioimmunoprecipitation assay buffer
ROS: reactive oxygen species
TPP: triphenylphosphonium
USD: United States dollar

## SUPPORTING INFORMATION

**Supporting Information Figure S1.**
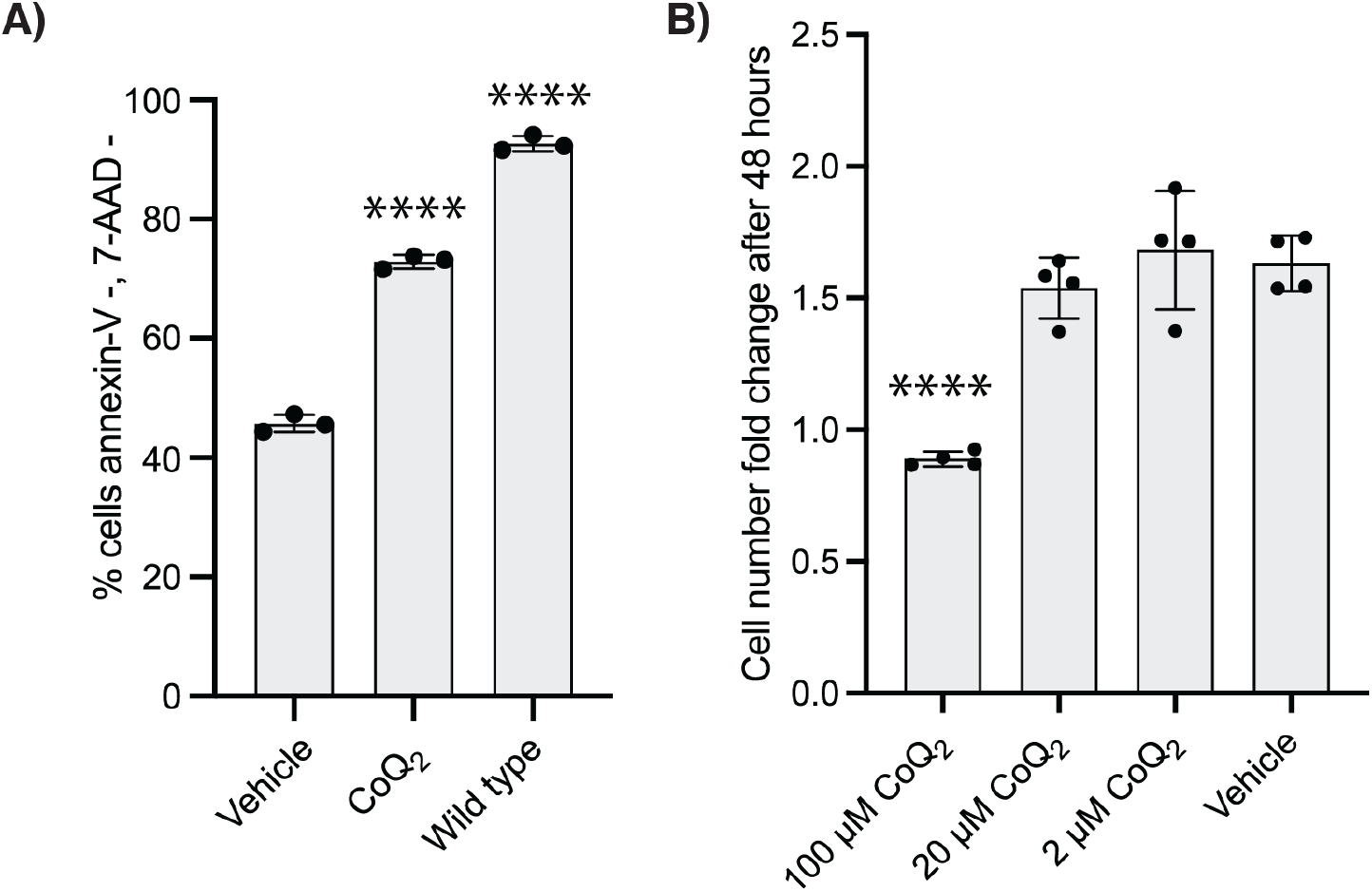
CoQ_2_ can support OxPhos but has inherent toxicity. **a** Percent non-apoptotic *COQ2*^*-/-*^ and wild-type HepG2 cells after 48 hours of incubation in galactose media supplemented with 20 µM CoQ_2_ or vehicle (isopropanol). Cells were pre-treated for 24 hours with CoQ_2_ to ensure sufficient uptake. Non-apoptotic cells defined as not staining for AAD-7 or annexin-V. *n* = three independent technical replicates, error bars indicate standard deviation. **b** Cell number fold change in wild-type HepG2 cells after 48 hours of incubation with indicated concentrations of CoQ_2_. *n* = four independent technical replicates, error bars indicate standard error. *ns: not significant, * p < 0*.*05, ** p < 0*.*01, *** p < 0*.*001, p < 0*.*0001*

**Supporting Information Figure S2.**
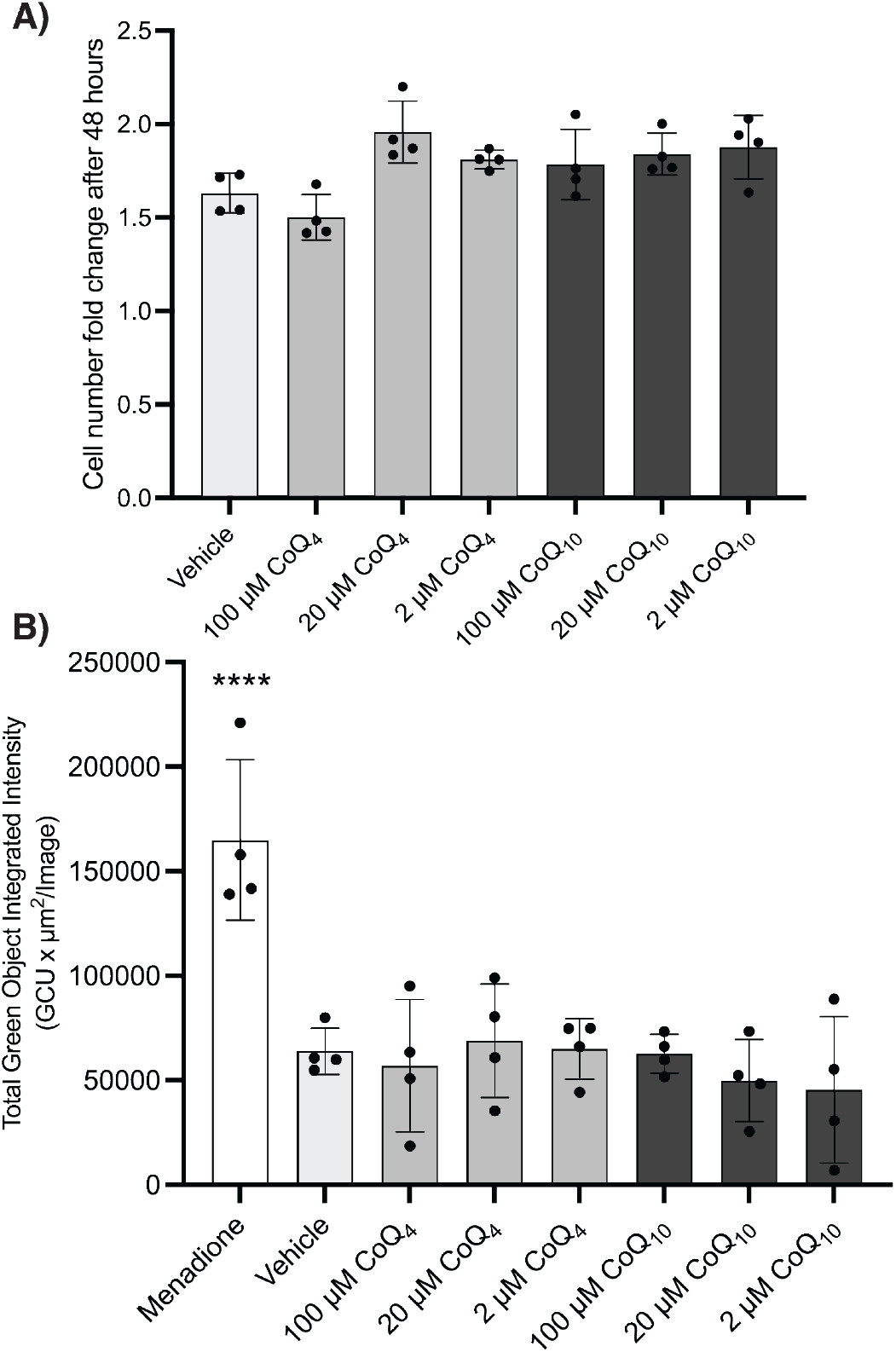
CoQ_4_ is not toxic at operative concentrations. **a** Cell number fold change in wild-type HepG2 cells after 48 hours of incubation with indicated concentrations of CoQ_4_ or CoQ_10_. **b** ROS production as visualized by green fluorescence from DCFDA in wild-type HepG2 cells after 24 hours of treatment with a positive control (25µM menadione) or indicated CoQ additives. *n* = four independent technical replicates, error bars indicate standard deviation. *ns: not significant, * p < 0*.*05, ** p < 0*.*01, *** p < 0*.*001, p < 0*.*0001*

**Supporting Information Figure S3.**
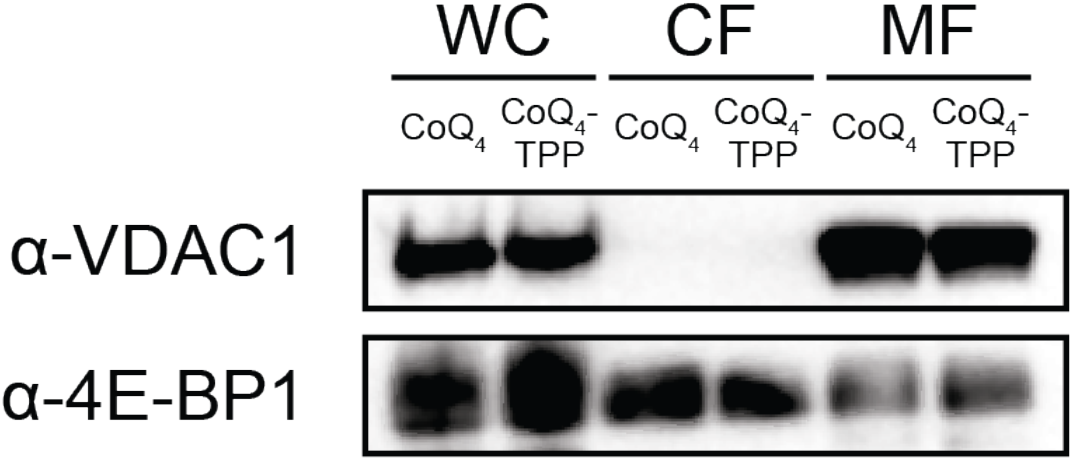
Crude mitochondria isolation blot. Enrichment of α-VDAC1 (mitochondrial marker) or α-4E-BP1 (cytoplasmic marker) in samples after mitochondrial preparation. WC – whole cell, CF – cytoplasmic fraction, MF – mitochondrial fraction.

**Supporting Information Figure S4.**
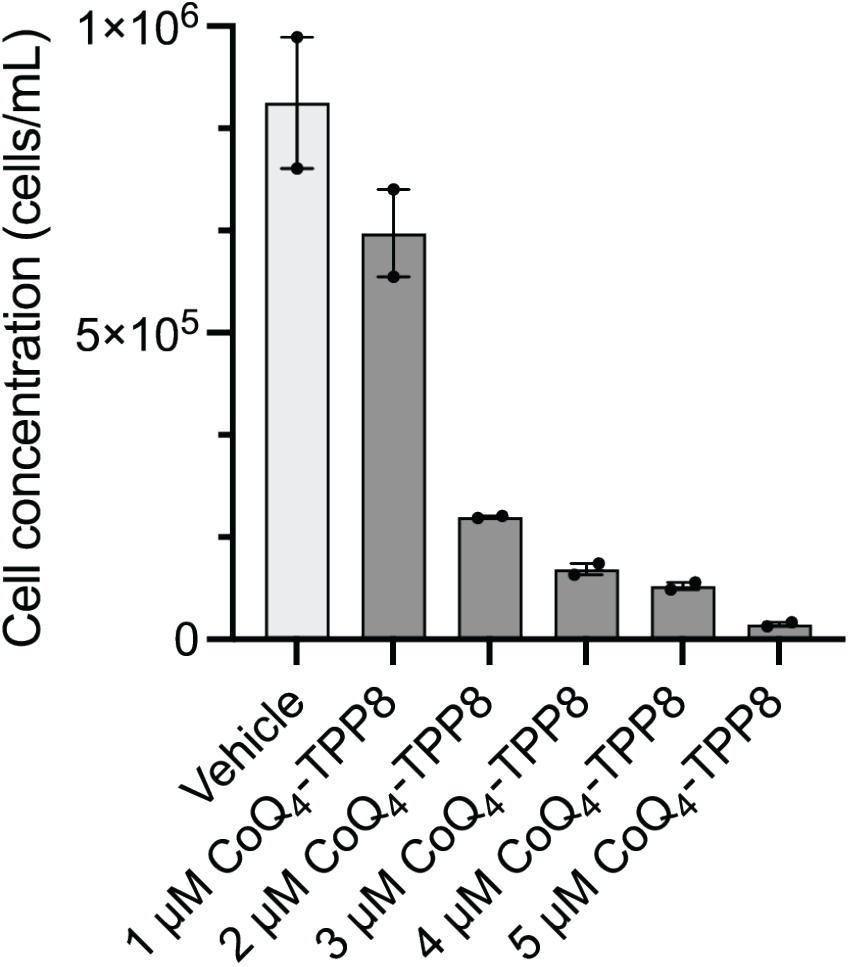
CoQ_4_-TPP compounds cause cell death. Cell concentration after 24 hours of incubation with various concentrations of CoQ_4_-TPP8. *n* = two independent technical replicates, error bars indicate standard deviation.

